# Unveiling the Developmental Dynamics and Functional Role of Odorant Receptor Co-receptor (*Orco*) in *Aedes albopictus*: A Novel Mechanism for Regulating Tuning Odorant Receptor Expression

**DOI:** 10.1101/2025.05.06.652105

**Authors:** Hui Yao, Qian Qi, Dan Gou, Simin Liang, Stephen T. Ferguson, Heng Zhang, Zi Ye, Feng Liu

## Abstract

As one of the most aggressive disease vectors, the Asian tiger mosquito *Aedes albopictus* relies heavily on its olfactory system to search for food in the larval stage, locate hosts after eclosion, and identify suitable oviposition sites after blood feeding. In mosquitoes and other insects, the olfactory system detects environmental odors primarily through a diverse repertoire of tuning odorant receptors (ORs), which require the highly conserved odorant receptor co-receptor (*Orco*) to function. While *Orco*’s role in enabling tuning receptor function is well established, its cellular localization patterns, developmental expression dynamics, and system-wide impact on olfactory physiology and behavior remain understudied in *Ae. albopictus*. To address this knowledge gap, we leveraged the Q-system to systematically characterize *Orco*-expressing neurons (ORNs) across embryonic, larval, pupal and adult stages of *Ae. albopictus*. We uncovered a dramatic reorganization of the olfactory system during metamorphosis: ORNs were observed as early as the embryonic stage and expanded during larval development before rapid degeneration and re-generation in the pupal stage resulting in the final population of adult ORNs. *Orco* expression in adults spanned the olfactory neurons of the antennae, labella, and maxillary palps in both male and female mosquitoes, consistent with its conserved peripheral distribution. To further investigate the functional implications of *Orco*, we generated *Orco* knockout mutants and strikingly discovered that *Orco* knockout mutants displayed significant widespread downregulation of tuning Ors, suggesting that *Orco* may influence OR expression or stability. Electrophysiological recordings confirmed significantly attenuated responses to human volatiles in *Orco* mutants, and behavioral assays demonstrated a marked decline in blood-feeding efficiency and elimination of host preference in females. Together, these findings reveal dynamic reorganization of ORNs during mosquito development and uncover the critical role of *Orco* in maintaining the integrity and function of the olfactory system, providing insights that may inform novel, next-generation vector control strategies.

## Introduction

The Asian tiger mosquito, *Aedes albopictus*, is a major vector of human disease, including dengue, chikungunya, Zika, and lymphatic filariasis and poses significant risks to global public health^1,2^. Dengue fever is one of the most severe tropical diseases transmitted by *Aedes* mosquitoes with models placing estimates around 390 million cases every year^3^ and 4 billion people at risk^4^. The threat of dengue fever continues to grow at an alarming rate, with the World Health Organization (WHO) reporting that from 2000-2019, there was a 10-fold increase in the number of documented cases being reported^5^. This is, in part, due to global climatic events such as the El Niño-Southern Oscillation phenomena^6^, increases in temperature and humidity associated with climate change, and population growth, leading to the expansion of suitable habitats for *Aedes* mosquitoes^7^. Importantly, *Ae. albopictus* has replaced *Ae. aegypti* as the primary vector of dengue virus in several countries, such as China and Japan, with its invasive range expanding at a far greater rate than *Ae. Aegypti* due to its superior ability to compete for breeding sites^8,9^. In light of the growing public health threat posed by *Ae. albopictus* and its rapidly expanding geographic range, a comprehensive understanding of the olfactory system, an essential driver of host-seeking behavior, is critical for developing timely and effective vector management strategies.

*Ae. albopictus* and other mosquitoes rely on their olfactory system for essential behaviors such as host-seeking and oviposition. The primary olfactory appendages in mosquitoes include the antennae, maxillary palps, and labella. Along these appendages are small sensory hairs known as sensilla that house sensory neurons responsible for the detection of various chemical, thermal, and mechanical stimuli^10^.Within the chemosensory neurons, insects express one or more major classes of chemoreceptor: odorant receptors (ORs), ionotropic receptors (IRs), and gustatory receptors (GRs)^11,12,13,14^. Chief among these are the olfactory receptor neurons (ORNs) which are responsible for detecting volatile odorants and pheromones, such as those associated with human sweat, and guiding mosquitoes towards potential hosts^15,16^.

Each of these neurons express one of many potential tuning odorant receptors (ORs) which convey ligand-specifity, alongside a highly conserved obligate co-receptor (*Orco*)^17^. Together, these protein subunits form a heterotetrameric (1:3 OR:Orco) ligand-gated ion channel on the membrane of the ORN^18,19,20,21^. Upon binding their cognate ligand, the channel undergoes a conformation change that leads to the opening of the central pore, allowing the influx of ions that depolarize the neuron and generate an action potential^20^. This electrical signal is transmitted to the brain, where it is processed to guide behaviors such as navigating towards hosts and selecting egg-laying sites.

Among the various molecular components involved in OR-mediated olfactory signal transduction, *Orco* is of central importance, and its conservation across diverse insect species highlights its essential contribution to the evolution and function of the insect olfactory system. Indeed, only certain species from the most basal insect order, Archaeognatha, appears to lack this essential gene^22^. Moreover, numerous studies across diverse insect lineages, including mosquitoes, have consistently demonstrated that the loss of *Orco* function dramatically impairs olfactory sensitivity and physiology, leading to aberrant behaviors. In *Drosophila*, *Orco* knockout significantly impairs olfactory responses to a broad range of general odorants and disrupts behavior in valence bioassays in larvae and adults^23^. In ants, both genetic knockout of *Orco* and pharmacological modulation of Orco protein significantly alter olfactory sensivity to hyodrocarbons and other important social chemical cues leading to the loss of social behaviors^24,25,26^.In *Aedes* and *Anopheles*, *Orco* knockout impairs the mosquitoes’ sensitivity to both human- and non-human-derived odor cues resulting in diminished attraction to human hosts and oviposition sites^27,28^. Taken together, these studies reinforce *Orco*’s central role in olfactory signaling and make it a prime target for developing vector control strategies.

While much is known about *Orco*’s fundamental role as an obligate co-receptor, studies continue to uncover its broader involvement in the development and maturation of olfactory systems, revealing new insights into how *Orco* functions beyond its role in sensory signal transduction. The functional role of *Orco* was first described in *Drosophila* studies, where its absence not only reduced olfactory sensitivity, but also severely disrupted the localization of ORs in the dendritic membranes of larval dorsal organs and adult antennae^17,23^. In *Orco*-deficient mutants, ORs were mislocalized to the cell bodies of ORNs instead of dendritic membranes, confirming *Orco*’s critical role in the proper localization and stabilization of ORs ^23^. Curiously, *Orco* seemed to have acquired novel functionality in different insect species over evolutionary time. In *Drosophila*, for example, *Orco* knockout leads to degeneration of olfactory neurons in both the antennal lobe and maxillary palps post-eclosion^29,30^; however, it does not lead to gross developmental or structural defects^23^. In Hymenopteran ants and honeybees, however, *Orco* mutants ORNs undergo massive apoptotic events during pupal development and display profound anatomical changes in the antennal lobe, including reduced antennal lobe volume and glomeruli number^24,25,31,32^. Similarly, *Orco* mutagenesis also leads to a reduction in the volume of pheromone-sensing macroglomeruli in male hawkmoths (*Manduca sexta*)^33^. These observations can be partially explained by differences in developmental timing. In *Drosophila*, the patterning of projection neurons in the antennal lobe glomeruli precedes the development of the ORNs, whereas in ants and moths, ORNs are formed before the glomeruli and ablation of these tissues leads to developmental defects in the antennal lobe^34,35^. By contrast, studies have thus far not reported ORN loss or major antennal lobe deficits in *Orco* knockout mosquitoes^27,28^, but emerging evidence suggests that *Orco* may play an important role in regulating OR transcription^36^ Together, these studies underscore the broad significance of *Orco* in the maintenance and function of the olfactory system in addition to its diverse roles in development across insect species. Furthermore, these findings emphasize the need to extend research efforts beyond model organisms like *Drosophila*, as exploring *Orco* function in lesser-studies species such as *Ae. albopictus* may reveal novel roles with potential implications for vector control strategies.

From these observations, we hypothesize that *Orco* plays a fundamental role in the development patterning of its peripheral olfactory system and the olfactory mechanisms underlying its host-seeking and blood-feeding behavior. To address this hypothesis, we performed a comprehensive characterization of *Orco*-expressing neurons across the developmental stages of *Ae. albopictus* mosquitoes. To elucidate the spatiotemporal expression patterns and function of *Orco^+^* sensory neurons, we employed a combination of advanced genetic and electrophysiological techniques. Specifically, we used the Q system^37^ to label *Orco*-expressing neurons via GFP, and integrated homology assisted CRISPR/Cas9 knockin (HACK) and PiggyBac transposon-mediated genomic integration to generate the *AalbOrco-QF2* driver line and *AalbQUAS-mCD8:GFP* reporter line. In parallel, we used RNA-sequencing, electroantennography (EAG), and single sensillum recording (SSR) to assess the impact of *Orco* homozygous knockout, while behavioral assays were used to examine the effects on blood-feeding efficiency and host preference. This study demonstrated a novel role of *Orco* in regulating OR transcription in mosquitoes and uncovered the dynamic organization of olfactory sensory neurons during *Ae. albopictus* development, particularly highlighting the remodeling of *Orco^+^* sensory neurons during the pupal stage. These findings demonstrated the critical role of *Orco* in chemical perception and host-seeking behavior in *Ae. albopictus*, providing a theoretical foundation for developing novel olfactory-based mosquito control strategies

## Result

### 1. Targeted CRISPR knockin of *AalbOrco*

To first characterize the spatial expression patterns of *Orco* in *Ae. albopictus* (Aalb*Orco*), we employed the binary expression Q system which enables precise visualization of gene expression through fluorescent signal reporting^37^ and has been successfully implemented in *Anopheles gambiae, Ae. aegypti* and *An. coluzzii*^38,39,40^.This system crosses an *AalbOrco promoter-QF2* (*AalbOrco-QF2*) driver line with a QUAS-GFP effector line to promote GFP expression in Aalb*Orco*-expressing cells. Here, we generated the driver line using the homology-assisted CRISPR/Cas9 knock-in (HACK) method^41^, targeting the fourth coding exon of *AalbOrco* by using guide RNA to insert a *T2A-QF2-3xP3-DsRed* cassette with a fluorescence red eye selection marker (Figure 1A, B, E). Successful integration was confirmed by genomic PCR amplification of the target locus (Figure 1C). Concurrently, we established the QUAS effector line using the piggyBac transposon system. This effector line carries the QUAS-mCD8:GFP-*3xP3-ECFP* reporter cassette with a cyan eye selection marker inserted into any TTAA site of the genome (Figure 1D, F). The *AalbOrco^+/DsRed^* virgin female was then backcrossed for five generations to wild-type males to minimize genetic variability and establish a stable transgenic mosquito line.

**Figure 1.**
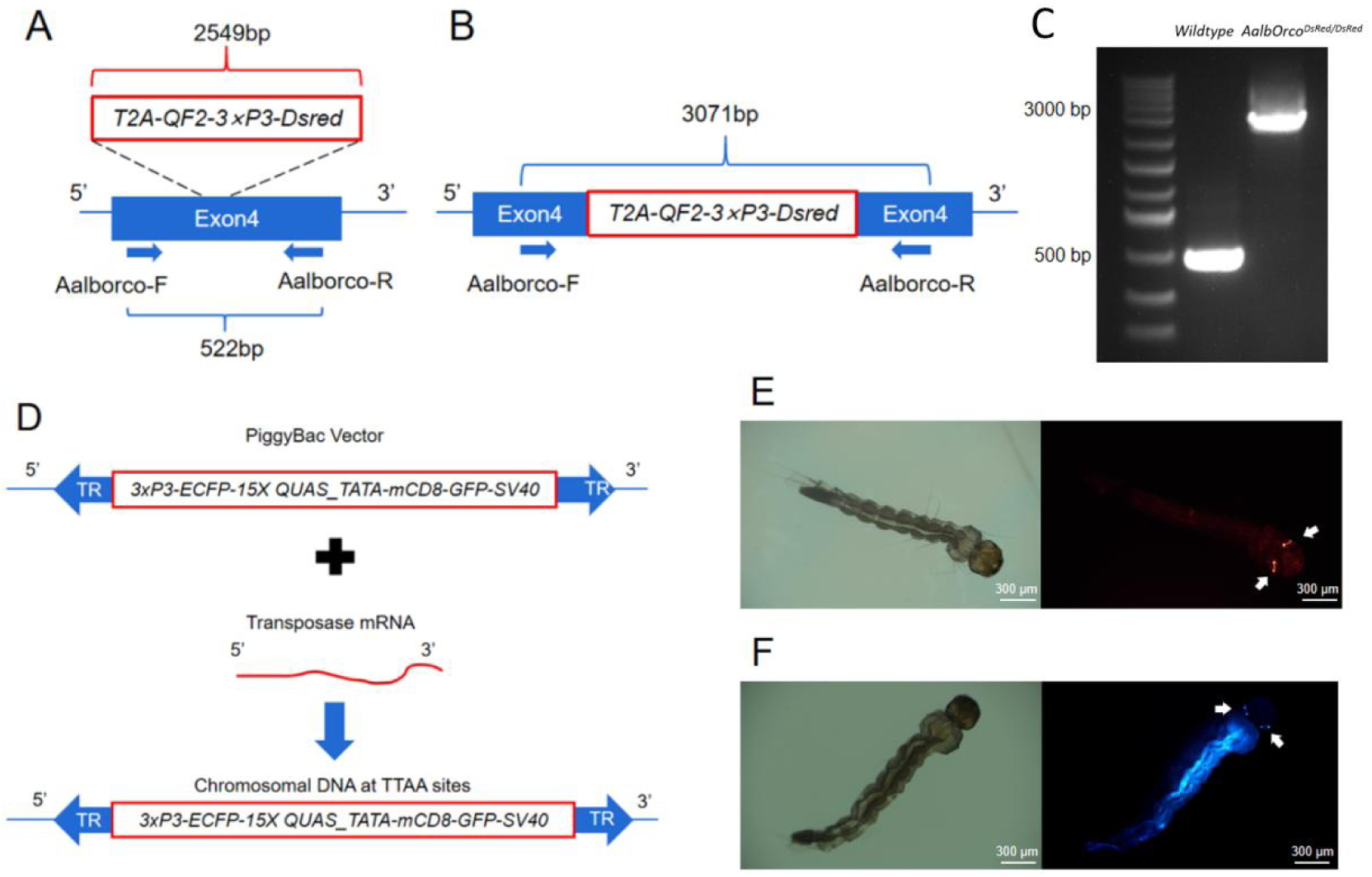
CRISPR-mediated construction of *AalbOrco-QF2* driver and AalbQUAS-mCD8:GFP reporter lines. (A) Homology-assisted CRISPR knock-in strategy. The *T2A*-*QF2-3xP3-DsRed* element (2549-bp) was inserted into the fourth exon of *AalbOrco* via CRISPR-mediated homologous recombination. PCR validation utilized primers *AalbOrco*-F/R (Table S1) and produced a 552-bp wild-type (WT) amplicon and a 3,071-bp mutant-specific product in homozygous individuals. (B) Primers *AalbOrco*-F and *AalbOrco*-R were designed to amplify the region outside the homology arm. This ensured allele-specific amplification, as the WT genome produced only the 552bp fragment, while the knock-in allele generated the 3,071bp band. (C) Agarose gel electrophoresis successfully confirmed knock-in integration in transgenic mosquitoes. The 3,071-bp band exclusively appeared in homozygous *AalbOrco^DsRed/DsRed^* individuals, while WT controls showed only the 552-bp fragment. (D) The *15xQUAS-GFP* cassette (5, 447-bp) was inserted into random TTAA sites in the genome via PiggyBac system. (E) In *AalbOrco-QF2* driver lines, *DsRed* fluorescence localized specifically to the eyes (scale bar: 300 μm), indicating functional promoter activity. (F) *AalbQUAS-mCD8:GFP* reporter lines exhibited ECFP fluorescence in the eyes, demonstrating transposon-mediated cassette insertion (scale bar: 300 μm).

### 2. Expression of *AalbOrco* in the late embryo of *Ae. albopictus*

Mosquito embryogenesis undergoes three major developmental stages: the initial stage, characterized by limited bristle development and cellular division during germ band extension; the intermediate stage, marked by bristle formation, segmentation of the cephalic and thoracic regions, and the development of structures such as the respiratory siphon; and the final stage, where complete segmentation occurs and the chorion-breaking spike forms preparing for larval hatching^42^. While the mosquito egg remains relatively understudies relative to other developmental stages, previous studies have demonstrated that certain chemical cues, such as those found in yeast, trigger embryo hatching, suggesting a potential chemosensory capacity of mosquito embryos^43^. To investigate the expression of *AalbOrco* in the late embryo stage of *Ae. albopictus*, we dissected the embryo chorion with an insect pin and directly imaged the GFP-labeled sensory neurons in the cephalic region of the embryo. While no GFP-labelled tissues had been observed in the initial (12 hours post-egg laying) and intermediate (36 hours post-egg laying) stage of mosquito embryoes, we found *Orco*^+^ neurons in the cephalic section of late embryos (72 hours post-egg laying) in two prepared samples after multiple trials (Fig. 2). Interestingly, one sample only displayed single Orco+ neuron (Fig. 2A) while the other sample showed 3-4 GFP-labeled Orco+ neurons (Fig. 2B) in the antenna-like tissue, suggesting the heterogeneity of Orco exprepression among individual embryoes in the development of olfactory sensory neuron. This result demonstrated the initial expression of *Orco* in the late embryonic stage of *Ae. albopictus*, providing support for the olfactory capacity in early development.

**Figure 2.**
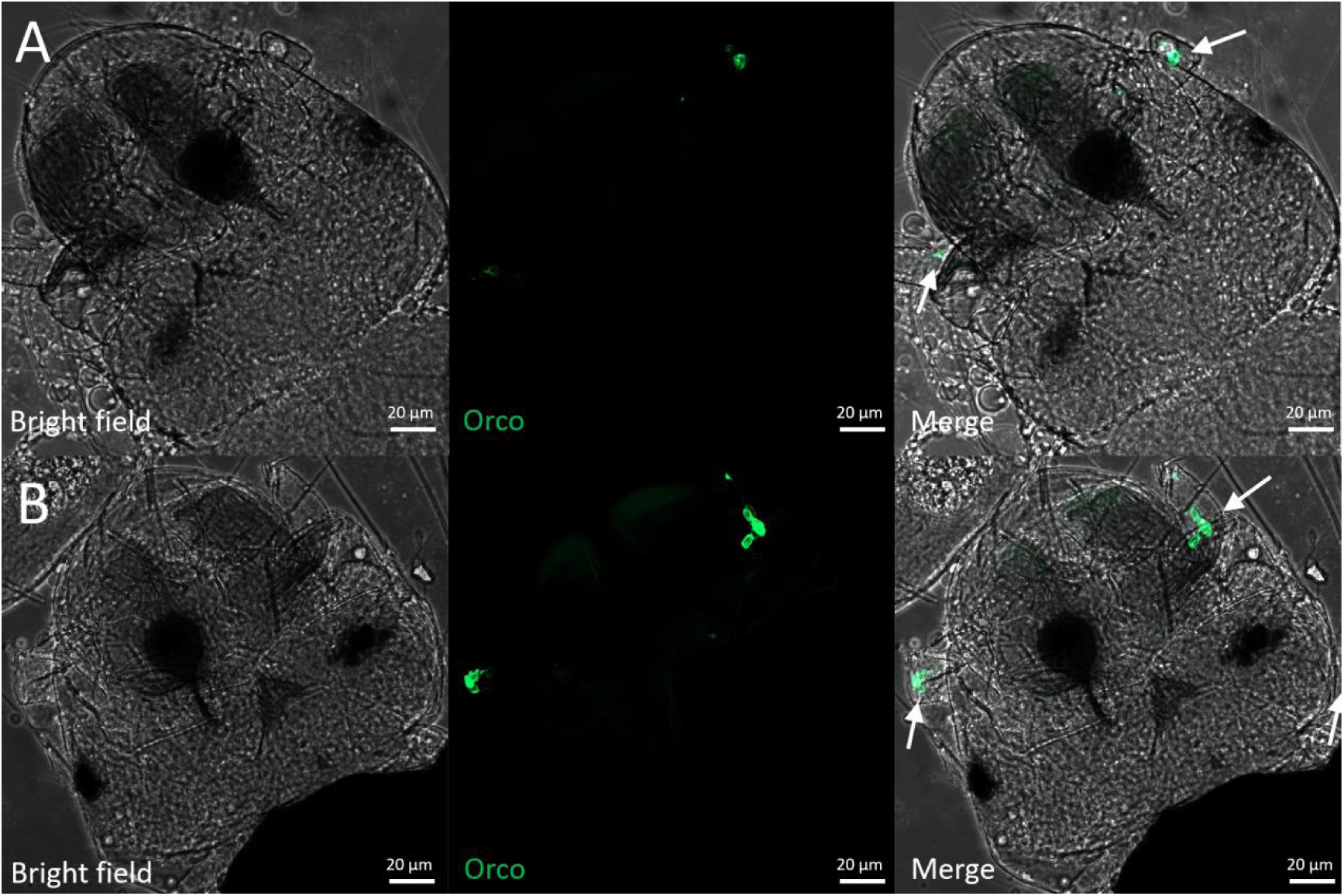
Localization of *Orco*-expressing neurons in the late embryo of *Ae. albopictus*. (A) & (B) Representative *Orco*-expressing neurons in the antenna-like tissue of the embryo at sample 1 (A) and sample 2 (B) 72 hours post-egg laying.

### 3. Expression pattern of *AalbOrco* in the larval antennae

Mosquito larvae possess a robust chemosensory capacity through a relatively simple olfactory system. To determine the spatial distribution of *AalbOrco* in the antennae of mosquito larvae, we systematically examined the progeny from crossing the *AalbOrco-QF2* driver lines with *AalbQUAS-mCD8:GFP* reporter lines. In these hybrids, GFP fluorescence specifically labeled neuronal dendrites, cell bodies, and axons in the larval sensory cone across all four instars (Fig. 3A-D). High-resolution confocal imaging showed that the number of *AalbOrco-*labelled olfactory neurons increased during larval development (N_L1_=5.25±1.40 (n=8), N_L2_=7±1.10 (n=6), N_L3_=11±0.89 (n=6), N_L4_=11.93±1.49 (n=14); Fig. 3E). Notably, the number of *Orco*-expressing neurons increased the most during the transition from 2^nd^ to 3^rd^ instar larva, which correlates with the considerable feeding needs and highly active food-searching behavior of 3^rd^ instar larva.

**Figure 3.**
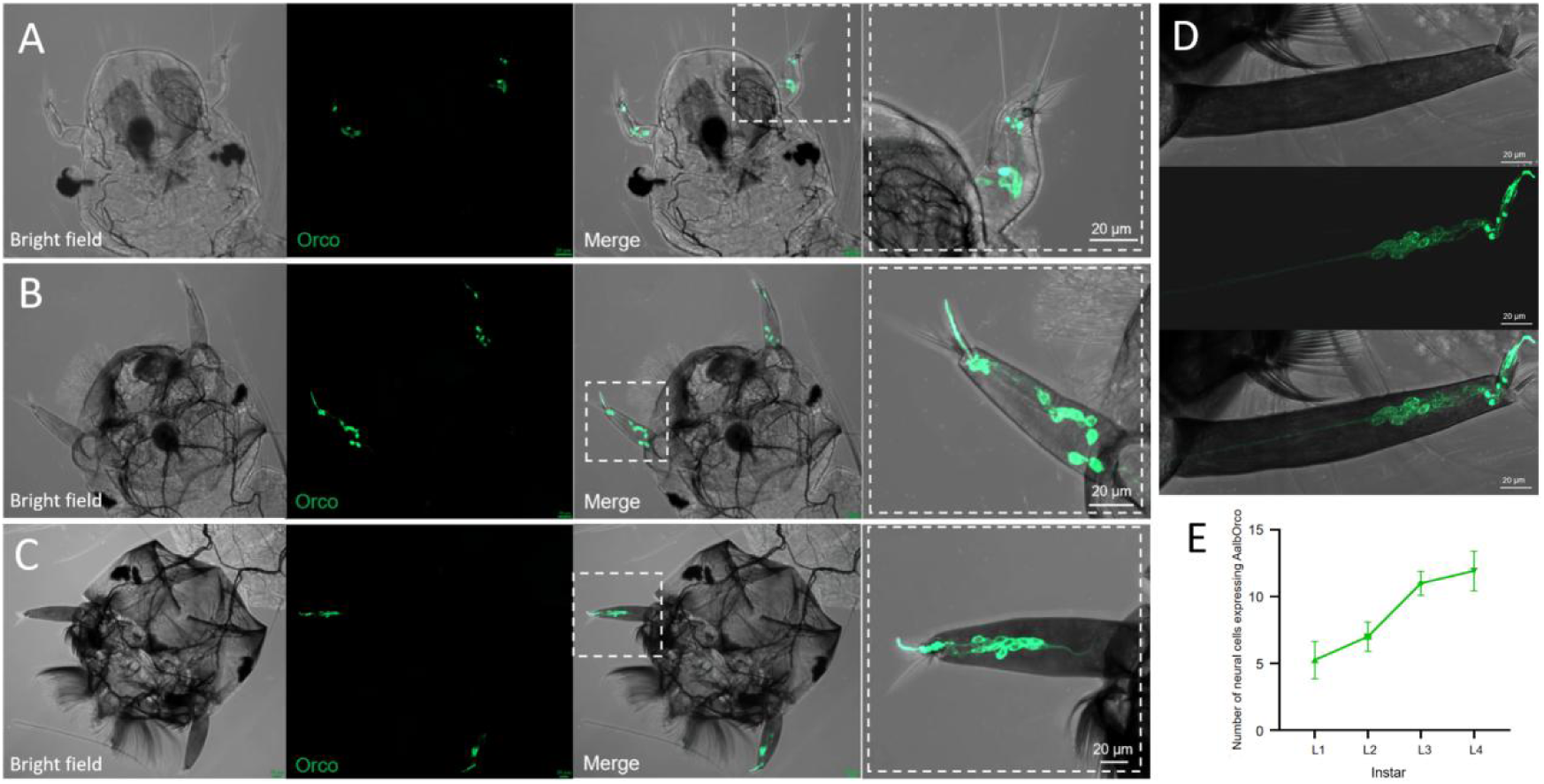
Localization of *Orco*-expressing neurons in the larval antenna of *Ae. albopictus*. (A-D) Maximum intensity projections of z-stack images showing GFP-labeled neurons in the antennae of 1-4 instar larvae. Scale bar: 20 μm. (E) Quantitative analysis of *Orco*^+^ neurons in 1-4 instar larvae. The number of *Orco*^+^ neurons increases with developmental stage, N_L1_=5.25±1.40(n=8); N_L2_=7±1.10(n=6); N_L3_=11±0.89(n=6); N_L4_=11.93±1.49(n=14).

### 4. Neurological remodeling in the pupa stage of *Ae. albopictus*

To elucidate the neuroanatomical changes during metamorphic development of *Ae. albopcitus*, we performed confocal microscopic analysis of the remodeling process of *Orco*^+^ neurons across the late 4^th^ instar larvae and early pupal stage. Using high-resolution labeling of *Orco*-expressing neurons, we revealed distinct stages of neurodegeneration and regeneration. For example, the antennae of early 4th instar (24 h post exuviation) larvae exhibited clear *Orco*^+^ neuronal cell bodies with nerve fibers concentrated into bundles (Fig. 4A). However, these *Orco*^+^ neurons and their associated dendrites began to degenerate in the wandering 4^th^ instar larvae (48 h post exuviation) (Fig. 4B). Moreover, disintegration of *Orco*^+^ neurons became even more pronounced after 96 hours where the neuron soma appeared to shrink, compress, and migrated towards the distal end of the antenna, and the neuron membrane became blurred, indicating active cell apoptosis (Fig. 4C). Notably, dramatic neuronal apoptosis in pupal antennae was observed after 3 hours of pupation with only residual GFP-labelled *Orco*-expressing cells in the tip of the antenna (Fig. 4D). In contrast to a limited number of *Orco*^+^ neurons in the larval antenna, numerous adult-like *Orco*^+^ neuron soma formed around 12 hours after pupation (Fig. 4E) and an almost complete adult antenna with fully developed *Orco*^+^ neurons and olfactory sensillum presented after 24 hours of pupation (Fig. 4F). These results demonstrate a rapid neural remodeling process in the pupal stage of Aedes mosquito, which underpins their adaption in developing ORNs as a strategy to prepare for the transition from a relatively simple aquatic habitat to a relatively more dynamic terrestrial environment.

**Figure 4.**
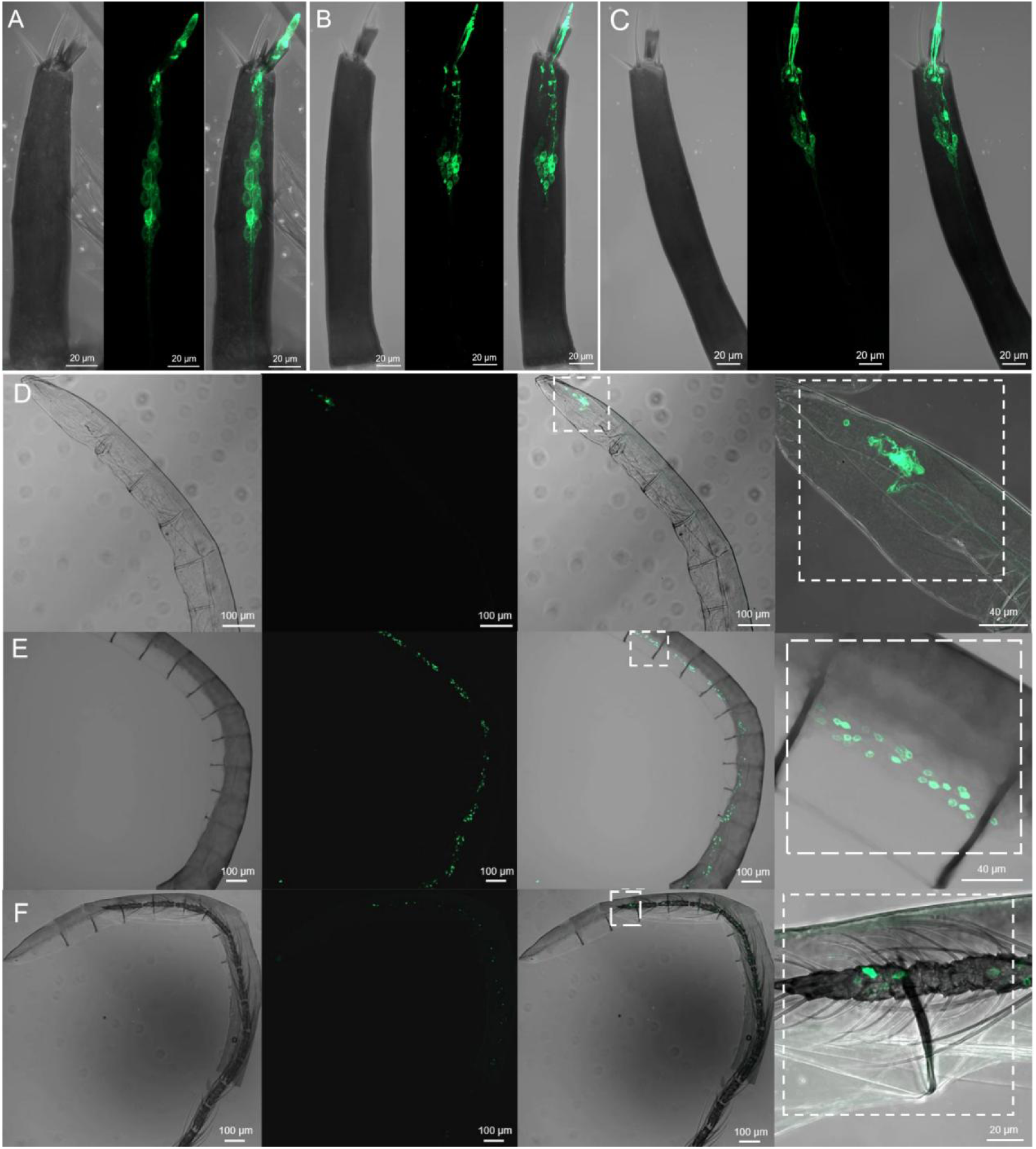
Remodeling of *Orco*-expressing neurons in the antenna of *Ae. albopictus* from early 4^th^ instar larva to early pupa stage. (A-C) Representative *Orco*-expressing neurons in the antennae of 4th instar larvae across different developmental time points ((A) 12h, (B) 48h, (C) 96h). (D-F) Representative *Orco*-expressing neuron in the antennae of female pupae at different time points ((D) 3h, (E) 12h, (F) 24h).

### 5. Expression pattern of *AalbOrco* in adult mosquito *Ae. albopictus*

To systematically investigate the spatial and sex-specific expression pattern of *AalbOrco* in *Ae. albopictus* adults, progeny derived from crosses between parental driver and effector lines were used for the whole-mount examination. Transgenically driven GFP expression was robust in ORNs of antennae, labella and maxillary palps. Expression patterns in the adult antennae were sexually dimorphic: female mosquitoes exhibited strong labeling across all 13 antennal segments, while male individuals displayed restricted expression on only the distal two segments (Fig. 5A, B). This sexually dimorphic pattern aligns with similar findings in *An. gambiae*^38^. *AalbOrco* expression in the labella was exclusively localized to ORNs with short dendrites associated with olfactory T2 sensilla as opposed to gustatory T1 sensilla (Fig.. 5C), consistent with reports in *Ae. aegypti*^44^ and *An. coluzzii*^45^. Despite pronounced morphological divergence in maxillary palp segmentation between female and male mosquitoes (males have four segments while females only have three), the GFP-labelled *Orco*-expressing neurons demonstrated striking conservation with two neurons housed in each capitate peg sensillum in both male and female maxillary palp (Fig. 4D, E), aligning with previous findings in both *An. coluzzii* and *Ae. aegypti*^44,45^.

**Figure 5.**
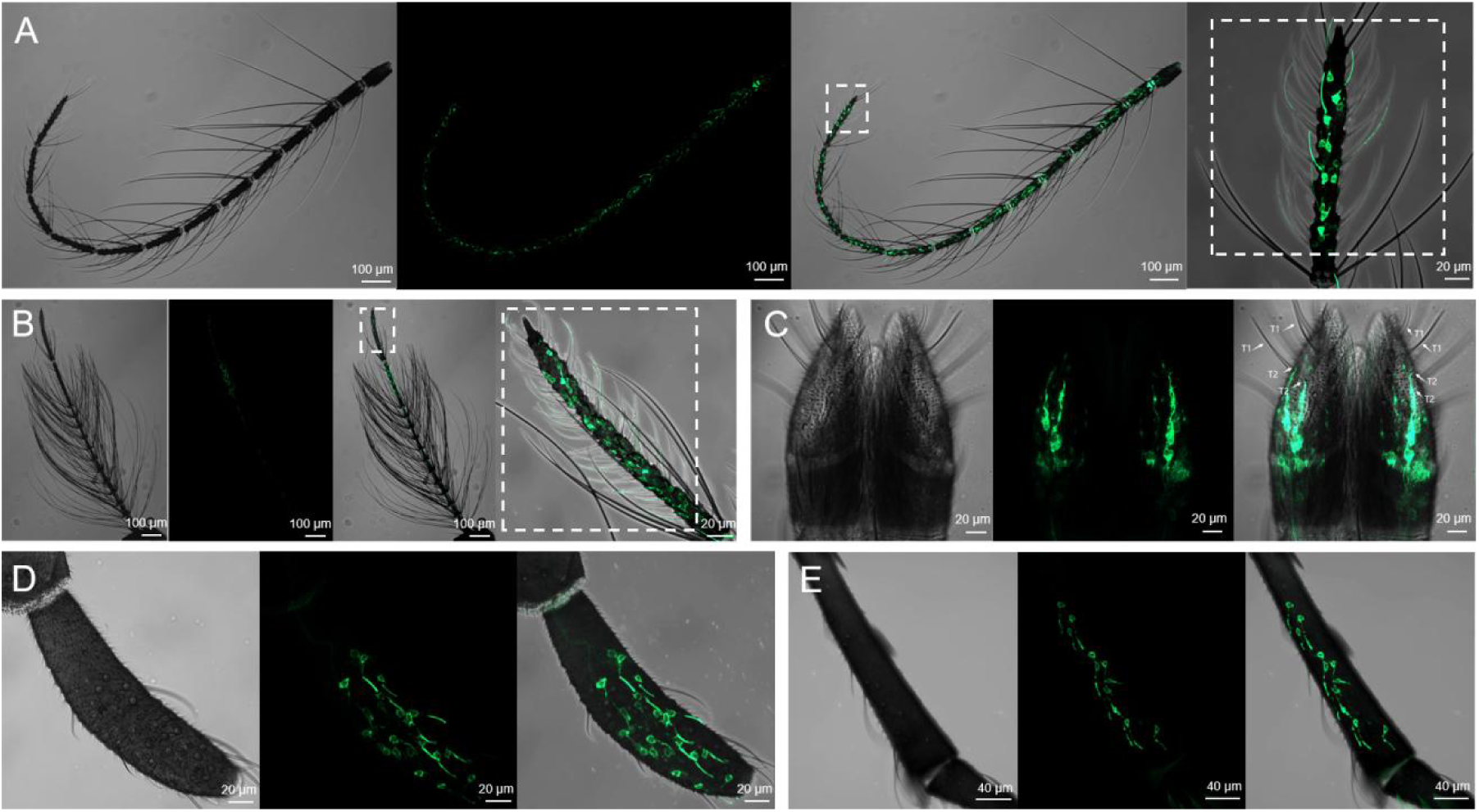
Localization of *Orco* in olfactory appendages of *Ae. albopictus*. (A-B) Representative confocal z-stack of *Orco*-expressing neurons in the antenae of female (A) and male (B) mosquitoes which exhibits sexual dimorphism with *Orco* expressed in all 13 segments of the female antennae while only in the distal two segments of the male antennae. (C) A representative confocal z-stack of a whole-mount female labellum showing *Orco* is expressed in T2 sensilla but not T1 sensillum (highlighted by arrows; scale bars, 10 μm). (D and E) Representative confocal z-stack of *Orco*-expressing neurons in the capitate peg sensilla of maxillary palp of both female (D) and male (E) mosquito.

### 6. *Orco* knockout affects expression of tuning receptors in *Ae. albopictus*

To investigate the impact of *Orco* mutations on tuning *Ors* expression in *Ae. albopictus*, we conducted transcriptomic analysis by antennal RNA-seq to compare expression of *Or* tuning receptors in female wild-type and *Orco^DsRed/DsRed^* homozygous mutants. As described in this study, as the T2A-QF2-3xP3-DsRed element was inserted in an exon of *Orco*, this directly disrupting its normal function, we self-crossed the heterozygous *Orco^+/DsRed^ Ae. albopictus* and then obtained the homozygous *Orco^DsRed/DsRed^* mosquito based on the strength of fluoresence in the mosquito eye. Antennal RNA extracted from the wild-type and *Orco^DsRed/DsRed^* mosquitoes four day post eclosion was submitted for next-generation transcriptional profiling, and receptor transcript abundance was quantified with software FeatureCounts (Dataset S1, “FPKM”). As expected, *Orco* expression is significantly reduced in *Orco^DsRed/DsRed^* mutants mosquitoes (Fig. 6A, B). The extremely modest number of residual *Orco* transcripts are most likely derived from the sequence in front of the double stranded break located in the fourth exon of the mutant mosquitos (Fig. 6B). More importantly, we found that the knockout of *Orco* leads to significantly reduced expression of many *Or* tuning receptors (Fig. 6A). The greatest reduction is seen for *Or52*, whose transcript in *Orco^DsRed/DsRed^* mutants is almost undetectable (Fig. 6C). The transcript abundance of many other tuning receptors was also significantly diminished, with less than 20% of WT level expression, including: OR7a (1%), OR13a (5%), OR132 (6%), OR56a (8%), OR43 (8%), OR6 (8%), OR10 (17%), OR122 (17%), OR84 (18%) and OR111 (19%) (Fig. 6C). Interestingly, 2 *Or* transcripts, OR115 and OR85, exhibited upregulation, with significant increases of 87% and 60%, respectively, relative to WT controls (Fig. 6F).

**Figure 6.**
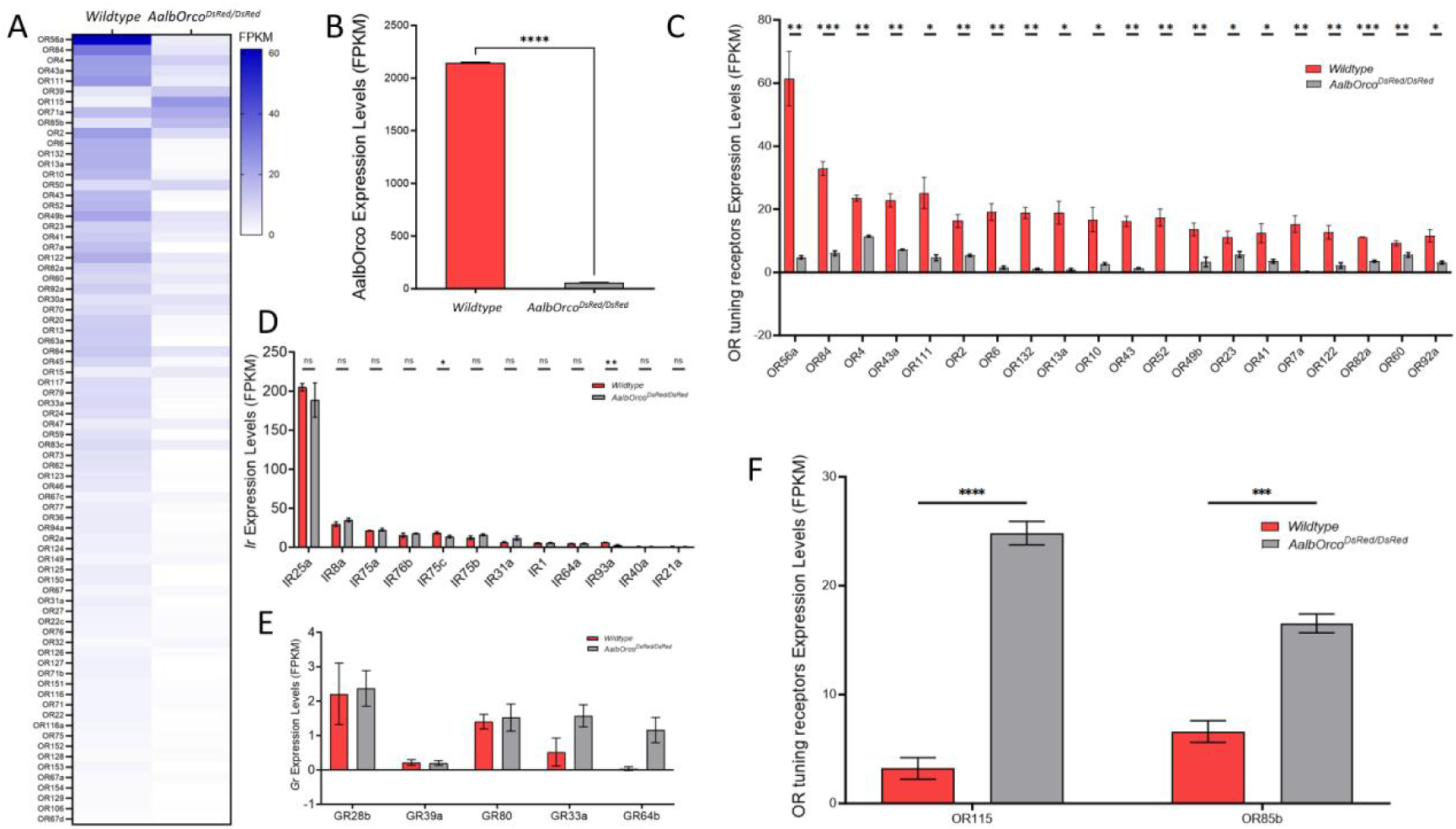
Comparison of expression of *Or* tuning receptors between the wild-type and *Orco^DsRed/DsRed^ homozygous mutant* mosquitoes. (A) Heatmap showing the average antennal expression in FPKM for each of the *Or* genes in WT and *Orco^DsRed/DsRed^* mosquitoes. (B) Transcriptional level of *Orco* gene in WT and *Orco^DsRed/DsRed^* mosquitoes. (C) Top 20 most highly expressed *Or* genes in WT were significantly reduced *Or* tuning receptors genes in *Orco^DsRed/DsRed^* mosquitoes. (D) Expression in FPKM for detected *Ir* tuning receptor and coreceptor genes in WT and *Orco^DsRed/DsRed^* mosquitoes. (E) Expression in FPKM for detected GR genes in WT and *Orco^DsRed/DsRed^* mosquitoes. (F) Expression of all the two upregulated *Or* tuning receptors genes in the *Orco^DsRed/DsRed^* mosquitoes. Unpaired Student’s *t*-test was applied in the statistical analysis, statistical significance is presented as P < 0.05 (*), P < 0.01 (**), P < 0.001 (***), P <0.0001(****) and P > 0.05(ns).

In contrast, no significant differences were observed in the transcript abundance of IRs in the antennae of *Orco^DsRed/DsRed^* mosquito (Fig. 6D). While GRs are generally considered absent from antennal expression, we nevertheless detected five *GR* transcripts of exceptionally low abundance in the antenna of both *Orco* homozygous mutants and wild-type mosquitoes, although these notably exhibited no significant difference between the two mosquito lines (Fig. 6E).

### 7. Reduced antennal responses of *Orco* knockout mosquito to human volatiles

To investigate the impact of *Orco* knockout on the compound-evoked olfactory responses of *Ae. albopictus*, we performed comparative electroantennogram (EAG) recordings between wild-type and *Orco^DsRed/DsRed^* homozygous mutant mosquitoes with a chemically diverse odorant panel (53 compounds across 9 chemical categories) which were selected according to their potent electrophysiological effects in other mosquito species^28,46,47^(Table S1). As expected, wild-type mosquitoes exhibited robust EAG responses across all of these chemical groups (Fig. 7A). In contrast, EAG responses of *AalbOrco* mutant mosquito to most odorants, particularly those in the categories of alcohol, aldehydes, ketones, and esters, were dramatically reduced (Fig. 7B). As expected, no significant differences were observed between *Orco^DsRed/DsRed^* and wild-type in response to three acids, valeric acid, hexanoic acid, and butanoic-acid, which are often considered to be detected by IRs, suggesting the *Orco/Or* complex is not involved in the perception of these acidic compounds. These results provide strong evidence that *Orco* is necessary for olfactory chemosensation in *Ae. albopictus*.

**Figure 7.**
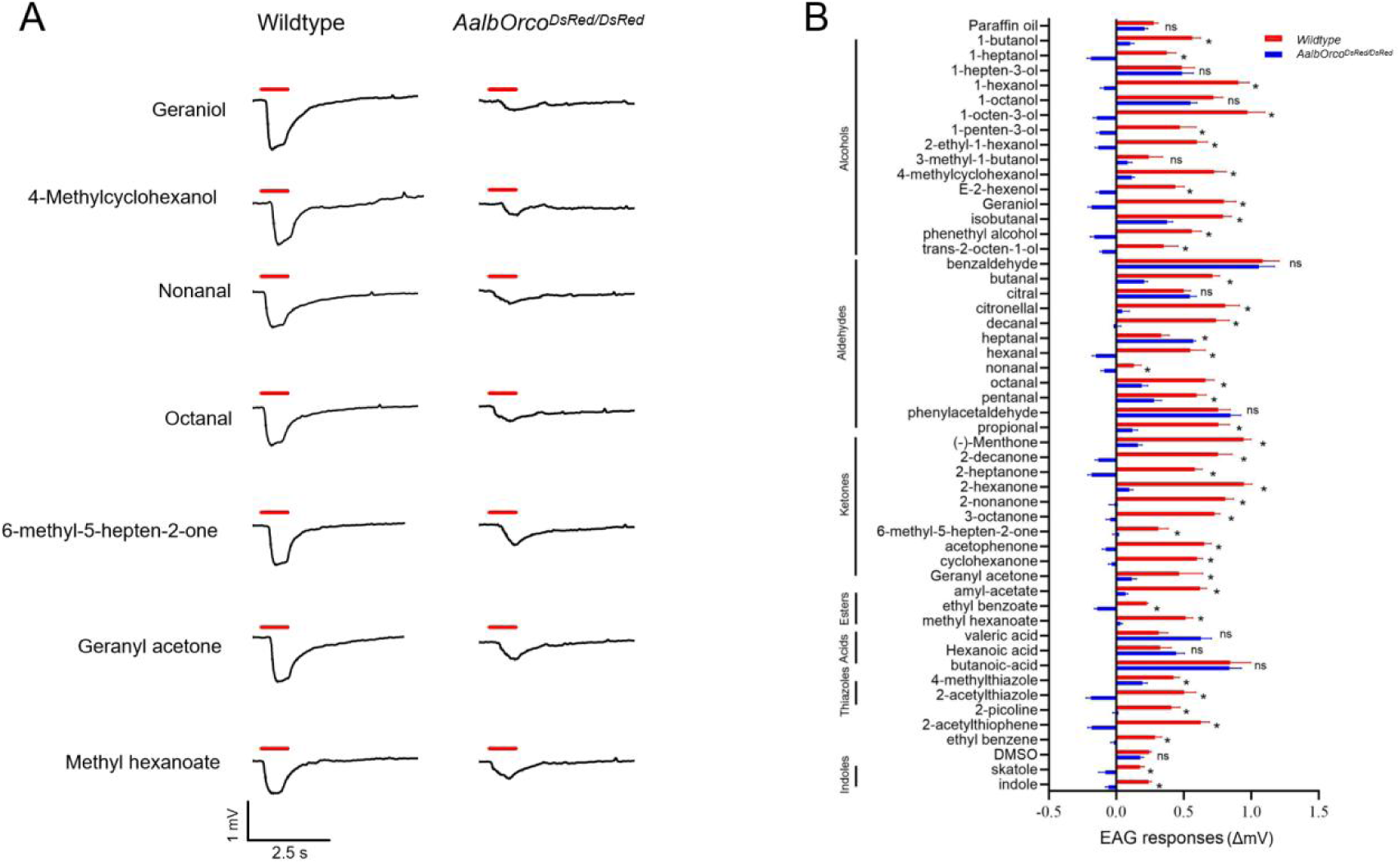
EAG responses of wild-type and *Orco^DsRed/DsRed^ Ae. albopictus* to a broad panel of human odorants. (A) Representative EAG response trace of wild-type and *Orco^DsRed/DsRed^* mosquito to different odorants; (B) Comparison of EAG responses of wild-type and *Orco^DsRed/DsRed^ Ae. albopictus* to 50 odorants in different chemical classes (n=7). EAG responses (ΔmV) for each odorant at a 10^−1^ dilution were normalized to the solvent control (paraffin oil and DMSO were used as solvents, among them, indole and methyl indole were dissolved in DMSO, while the other 48 compounds were dissolved in paraffin oil) by subtracting the solvent-induced EAG value (Table S2). Unpaired Student’s *t*-test was applied in the statistical analysis, with p ≥0.05 indicating no significance (ns), and P <0.05 (*) as significant differences.

### 8. Reduced Neuronal Responses in *Orco* mutant *Ae. albopictus*

Four morphological types of trichoid sensilla have been previously identified on the antennae of *Ae. albopictus* mosquitoes, namely long sharp tipped (LST), short sharp tipped (SST), short blunt tipped I (SBTI), and short blunt tipped II (LBTII)^48^. To investigate the effects of *Orco* knockout on chemical reception at the neuronal level, we performed SSR to compare the responses of four types of olfactory sensilla in wild-type and *Orco* mutants (*Orco^DsRed/DsRed^*) to a panel of 34 compounds in diverse chemical categories (Table S2). We found in female wild-type *Ae. albopictus*, sensilla SST and LBT exhibit broad spectrum neuronal responses to most compounds while LST and SBTII sensilla only respond to a limited number of chemicals (Fig. 8A). For example, the ’A’ neuron, as defined by its large spike amplitude, in SST (blue traces in Fig. 8C) displayed strong responses to 2,6-dimethylpyrazine (124 ± 54 spikes/s), 4,5-dimethylthiazole (120 ± 72 spikes/s), acetophenone (88 ± 44 spikes/s), and 1-pentanol (82 ± 62 spikes/s). Meanwhile, SBTI was very sensitive to terpenoid volatiles fenchone (82 ± 20 spikes/s) and camphor (72 ± 26 spikes/s) (Fig. 7C, left panel). However, in the *Orco^DsRed/DsRed^* mosquito, the trichoid sensilla, with the exception of SBTII, all exhibited loss of responsiveness to most compounds (Fig. 8B). Moreover, the background activity of sensory neurons in the sharp-tiped sensilla (SST and LST) were considerably diminished with very few residual spikes present (Fig. 8C, right panel). However, we did observe extensive spontaneous activity in the short blunt-tiped sensilla SBTI of the *Orco* mutant mosquito. Most interestingly, the short blunt-tiped SBTII sensilla are found to respond robustly to aldehydes-such as octanal, nonanal and decanal-in both wild-types and *Orco* mutants. The simple interpretation is that *Orco* is not involved in the sensation of aldehydes compounds in the SBTII sensillum of *Ae. albopictus*. Quantitative analysis on the neuronal response of *Orco* mutant mosquito revealed significant attenuation in SST sensitivity to 2,6-dimethylpyrazine, 4,5-dimethylthiazole, acetophenone, and 1-pentanol (Fig. 8D). In addition, SBTI also exhibited significantly diminished responses to fenchone and camphor (Fig.8E). This results demonstrate that *Orco*^+^ neurons housed in most trichoid sensilla are actively involved in the detection of a wide range of compounds.

**Figure 8.**
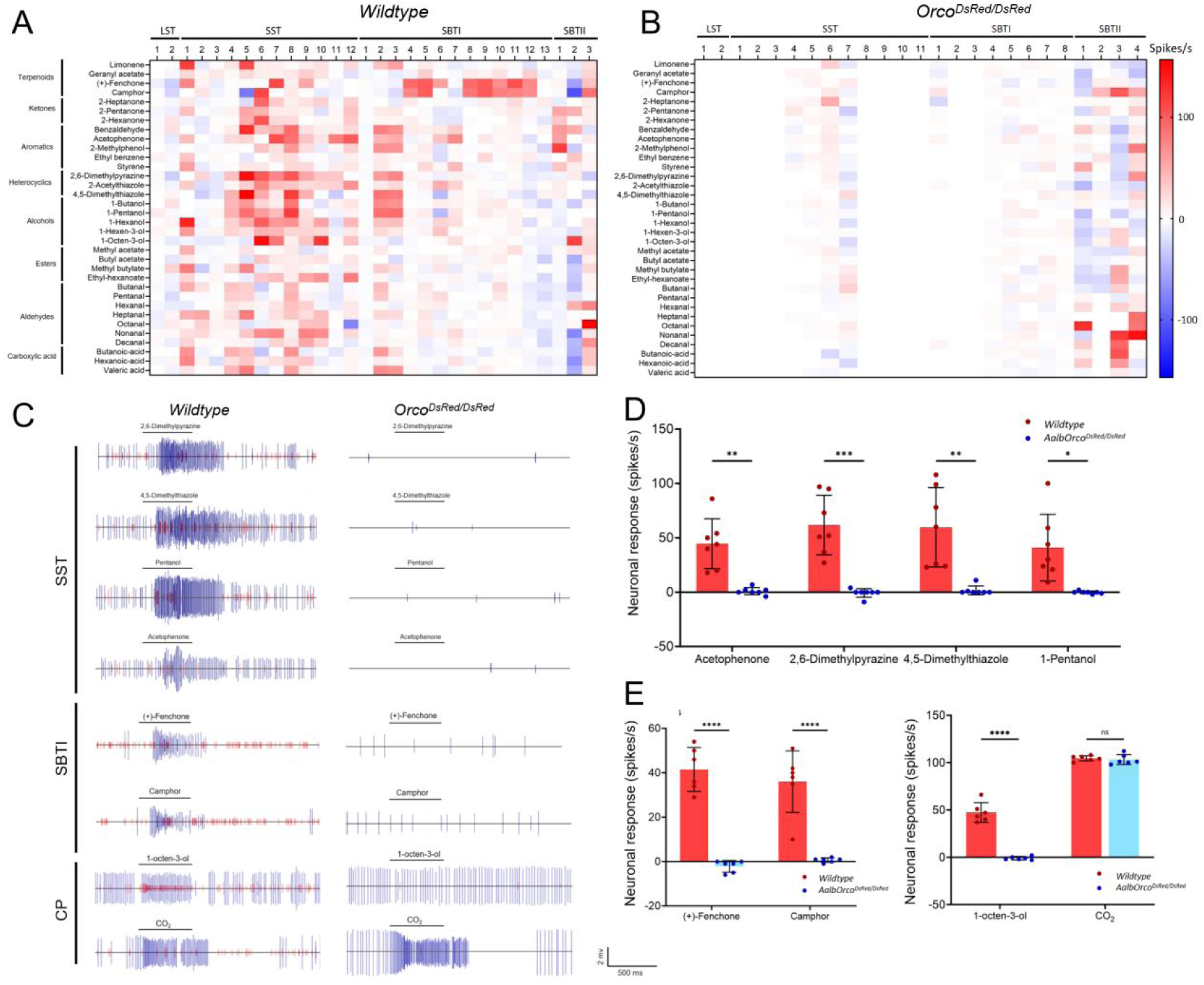
Responses of ORNs of different trichoid sensillum of wild-type and *Orco* mutant mosquito to odorants. (A) & (B) Heatmaps showing averaged female SSR responses to a 34 odorant panel of 33 (WT) and *Orco^DsRed/DsRed^* (28) individual sensilla randomly sampled across the antenna of wild-type (left) and *Orco^DsRed/DsRed^* (right) *Ae. albopictus*. (C) Representative response traces of SST and SBTI sensillum to selected chemical componds (diluted to 1% concentration in paraffin oil or DMSO). There are two neurons in most of the sensilla, in figure C, large amplitude (blue traces) represents the response of ’A’ neuron, small amplitude (red traces) represents the response of ’B’ neuron. (D) Averaged female SST poststimulus responses in WT and *Orco^DsRed/DsRed^* mosquitoes. Unpaired Student’s *t*-test suggests poststimulus responses to selected compounds in *Orco* mutants are significantly reduced compared to the wild type (n = 7). (E) Averaged female SBTI poststimulus responses in WT and *Orco^DsRed/DsRed^* mosquitoes. Unpaired Student’s *t*-test suggests poststimulus responses to fenchone and camphor in *Orco* mutants are significantly reduced than wild type (n=6). (F) Averaged female CP poststimulus responses in the the maxillary palp of WT and *Orco^DsRed/DsRed^* mosquitoes. Unpaired Student’s *t*-test suggests poststimulus responses to 1-octen-3-ol in *Orco* mutants are significantly reduced compared to the WT (n=6), while no significant differences were observed in poststimulus responses to CO_2_ between WT and *Orco* mutants (n=6). Statistical significance is presented as P < 0.05 (*), P < 0.01 (**), P < 0.001 (***), P <0.0001 (****).

Furthermore, we examined the odor-evoked responses of capitate peg (CP) sensillum which house three neurons, A neurons showing large spikes and B/C neurons displaying small spikes, on the maxillary palps of both wildtype and *Orco* mutant *Ae. albopictus*. The GR-expressing A neurons are responsible for detecting CO_2_ while the B/C neurons expressing the Orco/OR complex are extremly sensitive to one human sweat component: 1-octen-3-ol^27,49,50^. Our results revealed that in wildtype mosquitoes, the B/C neurons in CP sensilla of *Ae. albopictus* exhibited significantly higher responses to 1-octen-3-ol (47 ± 10 spikes/s) compared to *Orco* mutant’s, where the response to 1-octen-3-ol was almost completely abolished (Fig. 8F). In contrast, the CO_2_-detecting A neuron remained unaffected with both wildtype and *Orco* mutant mosquito displaying robust CO_2_-induced responses (Fig. 8F).

### 9. Impact of *Orco* knockout on the behavioral responses of mosquitoes

To further investigate the impact of *Orco^DsRed/DsRed^* mutation on blood-feeding and host preference, we conducted blood-feeding assays using mice as the blood source. We observed a significant decline in blood-feeding efficiency in *Orco^DsRed/DsRed^* mutants compared to the wildtype mosquitoes (P<0.05) (Fig. 9A and B). In parallel, a two-choice valence assay between volatile odors emitted from a human hand vs. that of a mouse was utilized to evaluate host preference of both wildtype and *Orco* knockout mosquitoes. As expected, wildtype mosquitoes displayed considerable preference towards the human hand (Fig. 9D). By contrast, host preference towards humans was significantly reduced and indeed entirely eliminated in *Orco^DsRed/DsRed^* mutant mosquitoes (P<0.0001) (Fig. 9C and D). Taken together, these findings make *Orco* a prime target for vector control strategies aimed at disrupting blood feeding and host preference in *Ae. albopictus*.

**Figure 9.**
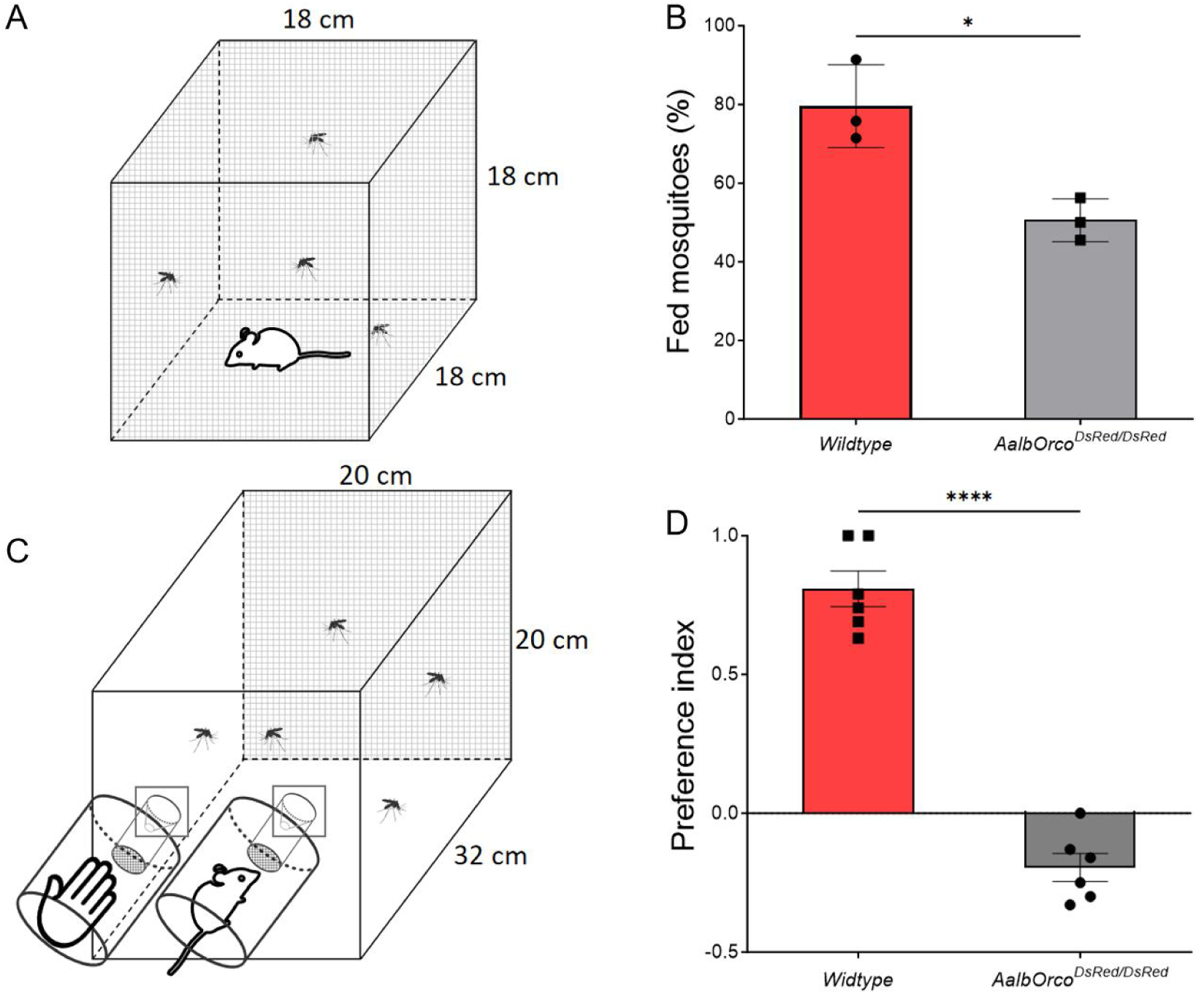
Behavioral assay. (A) Schematic picture of setting of blood-feeding assay. (B) Comparison of feeding success between *Orco^DsRed/DsRed^* and wild-type mosquitoes. The successful blood-feeding rate of the *Orco^DsRed/DsRed^* mosquito was significantly lower than that of wild-type mosquitoes on mice (P < 0.05, Unpaired Student’s *t*-test). (C) Schematic picture of setting of two-choice host preference assay. (D) Comparison of host preference between *Orco^DsRed/DsRed^* and wild-type mosquitoes. *Orco^DsRed/DsRed^* mutant mosquitoes showed preference for human hand (P < 0.0001, Unpaired Student’s *t*-test).

## Discussion

The emergence and global spread of the Asian tiger mosquito *Ae. albopictus* as a dominant arboviral vector represents a growing threat to public health, particularly given its expanding ecological range and capacity to transmit deadly diseases^1,2,7,8^. Despite the critical role of olfaction in the mosquito’s ability to locate hosts and reproduce, fundamental questions remain about how the olfactory system develops and functions in this species. In this study, we provide a systematic and comprehensive analysis of *Orco* expression, function, and regulatory impact across the life stages of *Ae.* albopictus, revealing an unexpectedly dynamic olfactory architecture and underscoring *Orco*’s multifaceted role in olfactory development, transcriptional regulation, and behavior.

Our study establishes that ORNs are present as early as the embryonic stage and expand throughout larval development, suggesting an early onset of olfactory capability that may contribute to environmental sensing, egg hatching, and larval foraging^43^. Here, we observed an average 11 *Orco*^+^ neurons in the third and fourth instar larval antenna, which is consistent with what has been reported in other mosquito species. For instance, the sensory cone in the larval antennae of *Ae. aegypti*, *An. coluzzii*, and *An. gambiae* (previously known as the “M” and “S” forms of the *An. gambiae* species complex)^51^ have all been reported to innervate 12–13 typical bipolar neurons that express Orco/OR complex and display robust chemosensory capacity^52,53,54,55^. Compared to several hundred *Orco^+^* neurons in the antenna of adult mosquitoes, the modest number of ORNs in the larval antenna may mirror the relatively less complex chemical environment of the aquatic system and their behavioral repertoire.

Most holometabolous insects undergo profound tissue reorganization during the process of complete metamorphosis, with some cells, tissues, and even entire organs being destroyed or completely reshaped, including the epidermis, nervous system, and muscles. This process involves programmed cell death and stem cell proliferation^56^. Strikingly, we uncovered a rapid and complete neuroanatomical reorganization of the peripheral olfactory system during metamorphosis, wherein larval ORNs undergo apoptotic degeneration and are replaced by a newly patterned adult population. The remodeling of peripheral neuronal system during metamorphosis mirrors observations in other holometabolous insects like the vinegar fly *D. melanogaster* and the hawkmoth *M. sexta*, whose peripheral nervous system also undergoes striking remodeling during the pupal stage^57,58^, but has not previously been demonstrated at cellular resolution in mosquitoes. Unlike other insects, however, our results suggest complete degeneration of larval sensory neuron in mosquitoes. By contrast, the axons of sensory neurons in the larval antennae of the red flour beetle *Tribolium castaneum* are found to be reused in the adult antennae^59^. Similarly, in *Drosophila*, certain type of peripheral sensory neurons, like C4da neurons, also found to be “recycled” during neuronal remodeling^60^. In *Drosophila*, the dendrites of C4da neurons are first severed from the soma and then degraded during metamorphosis while the soma and axons are kept intact. Subsequently, new dentritic arbors are regenerated on the soma in developing adult neurons. The partial survival of larval sensory neurons may be important for the assembly of the adult sensory system, and the absence of this process in *Ae. albopictus* suggest other mechanisms may be involved in mosquitoes. As it stands, the timing and precision of this transformation imply a tightly regulated olfactory developmental program tuned to the organism’s ecological transition from aquatic larva to terrestrial adult.

Functionally, *Orco* mutants exhibited a profound deficit in olfactory physiology. EAG and SSR demonstrated sharply reduced responses across multiple classes of odorants, including key human-derived volatiles such as 1-octen-3-ol and other aldehydes and ketones^16^. Indeed, the neuronal responses of most of the trichoid sensilla investigated in this study were almost completely abolished after *Orco* knockout. These deficits were sensory-specific, with GR- and IR-mediated responses largely preserved, reinforcing the selectivity of the Orco/OR pathway and corroborating earlier findings in *Anopheles* and *Aedes* species^27,28^. Behavioral assays further confirmed that *Orco* disruption abolishes host preference and significantly reduces blood-feeding success, firmly establishing *Orco* as a central node linking molecular, cellular, and behavioral levels of mosquito olfaction.

Curiously, we also observed abundant residual spikes in the sensilla of *Orco* mutants, particularly in the SbtI sensilla, which displayed a very high frequency of spontaneous sensory neuron activition. Our best explanation for these residual spikes comes from recent studies reporting non-canonical co-expression of chemoreceptor co-receptors (e.g., *Orco*, *Ir25a*, *Ir8a* and *Ir76b*) and/or tuning receptors in the olfactory sensory neurons of both *Drosophila* and mosquitoes^13,14,61,62,63^. The extensive presence of residual spikes in the SbtI sensillum of *Orco* mutant mosquito may therefore be the result of the co-expression of both Orco/OR and Irco/IR in the same neurons, which could potentially still be partially functional even though the *ORCO*/OR complex has been disrupted. Future studies on olfactory receptor organization at a single cell level in *Ae. albopictus* would give us a more definitive explanation of these residual spikes displayed after knocking out the *Orco* gene.

Beyond anatomical remodeling and physiology, our transcriptomic analysis revealed that *Orco* knockout in *Ae. albopictus* led to widespread downregulation and targeted upregulation of many tuning Ors, implicating *Orco* not only in ORN signaling but also in the maintenance of OR gene expression. These findings align with evidence suggesting a dual role for *Orco* in the transcriptional regulation of tuning *Ors* in *D. melanogaster* and *H. armigera*^64,65.^Together with a more recent study in *Ae. aegypti*^36^, our results demonstrate that this dual regulatory role of *Orco* in conserved in mosquitoes, as well, extending the functional landscape of *Orco* beyond its canonical role as merely a co-receptor^17,18^. The reduction in OR transcript levels suggests that *Orco* may influence receptor stability, trafficking, or transcriptional feedback, an area ripe for further investigation.

Taken together, this work provides the most detailed spatiotemporal characterization of *Orco* expression and function to date in a non-model mosquito species and establishes a new platform for studying olfactory system development. By integrating genetic tools (Q-system, HACK), transcriptomics, neurophysiology, and behavior, we demonstrate that *Orco* serves not only as a co-receptor but as a transcriptional stabilizer in *Ae. albopictus*. These findings provide a theoretical foundation for translational strategies aimed at disrupting mosquito host-seeking behaviors via genetic or chemical interference with *Orco* function. Given the expanding public health burden of *Ae. albopictus* and the increasing resistance to traditional insecticides, *Orco* and its associated pathways represent compelling targets for next-generation vector control.

## Materials and Methods

### Mosquito maintenance

Mosquito *Ae. albopictus* Foshan strain was a gift from Dr. Xiaoguang Chen from Southern Medical University, Guangzhou, China and reared as described^66^; and 5- to 7-day-old females that had not been blood fed were used for all experiments. All mosquito lines were reared in environmental chambers at 27°C and 75% relative humidity under a 12:12 light-dark cycle and supplied with 10% sucrose water in the Shenzhen Bay Laboratory Insectary.

### Electrophysiology

Single sensillum recordings (SSR) were conducted as described in Liu et al^67^. Female mosquitoes 4 days after eclosion were anaesthetized on ice for 2–3 min and mounted on a microscope slide (76 × 26 mm). The antennae were fixed using a double-sided tape to a cover slip resting on a small ball of dental wax to facilitate manipulation. Once mounted, the specimen was placed under a microscope (Eclipse FN1, Japan) and the antenna viewed at a high magnification (1000×). Two tungsten microelectrodes were sharpened in 10% KNO2 at 10 V. The reference electrode, which was connected to ground, was inserted into the compound eye of the mosquito and the other was connected to the preamplifier (10×, Syntech, Kirchzarten, Germany) and inserted into the shaft of an olfactory sensillum to complete the electrical circuit to extracellularly record ORN potentials^68^. Controlled manipulation of the electrodes was performed using a micromanipulator (Burleigh PCS-6000, CA). The preamplifier was connected to an analog-to-digital signal converter (IDAC-4, Syntech, Germany), which in turn was connected to a computer for signal recording and visualization in the software AutoSpike v5.1. Signals were recorded for 10 s starting 1 s before stimulation, and the action potentials were counted offline automatically with the AutoSpike software over a 500-ms period before and after stimulation. The spontaneous firing rates observed in the preceding 500 ms were subtracted from the total spike rates observed during the 500-ms stimulation, and counts were calculated in units of spikes/s.

Thirty-four compounds from different chemical classes were selected for electrophysiological recording of olfactory sensilla with various morphological shapes. Each compound was prepared in 100-fold dilution (v/v) with dimethyl sulfoxide (DMSO) or paraffin oil. For each dilution, a 10 μl portion was dispersed onto a filter paper strip (4 × 30 mm), which was then inserted into a Pasteur pipette to create the stimulus cartridge. A sample containing the solvent alone served as control. The airflow across the antennae was maintained constant at a 20 ml/s throughout the experiment. Purified and humidified air was delivered to the preparation through a glass tube (10-mm inner diameter) perforated by a small hole 10 cm away from the end of the tube, into which the tip of the Pasteur pipette could be inserted. The stimulus was delivered to the sensilla by inserting the tip of the stimulus cartridge into this hole and diverting a portion of the air stream (0.5 l/min) to flow through the stimulus cartridge for 500 ms using a stimulus controller (Syntech, Germany). The distance between the end of the glass tube and the antennae was ≤1 cm. The number of spikes/s was obtained by averaging the results for each sensillum/compound combination.

The electroantennogram procedure followed previously described protocols^69^ with minor modifications. Briefly, the head of an adult *Ae. albopictus* female was excised and mounted on an EAG platform equipped with two micromanipulators and a high-impedance AC/DC preamplifier (Syntech, Germany). Chlorinated silver wires in glass capillaries filled with 0.1% KCl and 0.5% polyvinylpyrrolidone were used for both reference and recording electrodes. One antenna with the tip cut was accommodated into the recording electrode. The airflow across the preparation was maintained constant at 20 ml/s to which a stimulus pulse of 2 ml/s was delivered for 500 ms. Any change in antennal deflection induced by the stimuli or control puffs was recorded for 10 s. All compounds were dissolved in DMSO or paraffin oil to make a test solution of 10-fold dilution. An aliquot (10 μl) of a tested compound was loaded onto a filter paper strip (4 × 30 mm), which was immediately inserted into a Pasteur pipette for evaporation. Solvent (paraffin oil) alone served as control. For each compound, EAG responses of 5 to 11 female mosquitoes were recorded.

### Chemicals

Compounds that were used in electrophysiological recordings are listed in Supplementary Table 1.

### sgRNA design and production

The procedure for single guide RNA (sgRNA) synthesis followed previously described methods^70^ with minor modifications. sgRNA (Supplementary Table 2) were designed for high efficiency by searching the sense and antisense strands of the *Orco* gene (AALFPA_042885) for the presence of protospacer-adjacent motifs (PAMs) with the sequence of NGG using CHOPCHOP^71^. sgRNAs were synthesized using the EnGen sgRNA Synthesis Kit (New England Biolabs) according to the manufacturer’s protocol using 300 ng of purified DNA template. Following in vitro transcription, the sgRNAs were purified using the Monarch RNA Cleanup Kit (New England Biolabs) and diluted to 1000 ng/μl in nuclease-free water and stored in aliquots at −80 °C.

Recombinant Cas9 protein from *Streptococcus pyogenes* was obtained commercially (TrueCut Cas9 Protein V2, Invitrogen by Thermo Fisher Scientifics) and diluted to 1000 ng/ul in nuclease-free water and stored in aliquots at −80 °C.

### CRISPR mediated microinjections

To generate the *AalbOrco-QF2* driver line for the Q system, *T2A-QF2-3xP3-DsRed* element was inserted into the *Orco* coding region through CRISPR-mediated homologous recombination. The homologous template (donor plasmid) that contains *T2A-QF2-3xP3-DsRed* element, which was amplified from the ppk301-T2A-QF2 HDR plasmid (Addgene plasmid# 130667)^39^, flanked by ∼1kb homologous arms was constructed using NEBuilder HiFi DNA Assembly kit (New England Biolabs). The left homologous arm was amplified with primer pair of OrcoleftarmFwd(GACGGCCAGTCAGGGGCGCTTCAAGTTAATAATTAAAAAA TAC) and OrcoleftarmRev(CTCTGCCCTCCGGACGGTAGGTGTCCAG). The right homologous arm was amplified with primer pair of OrcorightarmFwd (ATGTATCTTAACTCGGCTGCCCTGTTCC) and OrcorightarmRev (CAGCTATGACCGGCTCCGTGTGTAAGATCAC). The primer pair for comfirming the knockin element was OrcoQF2Fwd (CTACCGTCCGGAGGGCAGAGGAAGTCTTC) and OrcoQF2Rev(GCAGCCGAGTTAAGATACATTGATGAGTTTGGACAAAC). From the left homologous arm immediately preceding the T2A, 1 bp were removed to keep theT2A sequence in-frame. Red-eyed F1 mosquitoes were backcrossed for five generations and then crossed to the effector line to acquire progeny for *Orco* localization studies.

Embryonic collection and CRISPR microinjections were performed following the procedure described in Li et al^70^. Briefly, Aedes mosquitoes were blood-fed 4 days before egg collection. An ovicup filled with ddH_2_O and lined with filter paper was placed into a cage and female mosquitoes were allowed to lay eggs in the ovicup in the dark. After 30-60 min, the ovicup was taken out and unmelanized eggs were transferred onto a glass slide. The eggs were quickly aligned on a wet piece of filter paper. Aluminosilicate needles were pulled on a Sutter P-1000 needle puller and beveled using a KDG-03 beveler (Kewei, Wuhan). An Eppendorf Femtojet 4i was used for power injections under a compound microscope at 10× magnification (E5, Soptop, Ningbo). About 10 eggs were injected each time immediately after fresh eggs were collected. The concentration of components used in the study was as follows; Cas9 protein at 300 ng/μl, sgRNA at 40 ng/μl, donor plasmid 300 ng/μl. After injection, eggs were placed in a cup filled with water and allowed to hatch and develope into adults. The first generation (G0) of injected adults were separated based on sex and crossed to 5X wild-type counterparts. Their offspring (F1) were manually screened for DsRed-derived red eye fluorescence using an Compound Fluorescent Microscope (DP74, Olympus, Japan). Red-eyed F1 males were individually backcrossed to 5-fold wild-type females to establish a stable mutant line. DNA extraction was performed using FastPure Gel DNA Extraction (Vazyme Biotech, Nanjing) protocols and genomic DNA templates for PCR analyses of all individuals were performed (after mating) to validate the fluorescence marker insertion using primers that cover DSB sites (Table S1). PCR products were sequenced to confirm the accuracy of the genomic insertion. Heterozygous mutant lines were thereafter backcrossed to wild-type *Ae. albopictus* for at least five generations before self-crossing and the progenies were used for screening homozygous individuals according to their DsRed-derived red eye fluorescence intensity. Putative homozygous mutant individuals were mated to each other before being sacrificed for genomic DNA extraction and PCR analyses (as above) to confirm their homozygosity.

The reporter line (QUAS-mCD8:GFP) was generated by the plasmid pBAC-ECFP-15xQUAS_TATA-mCD8-GFP-sv40 (Addgene #104878)^38^ with a pBac helper plasmid. Similar to establishing the driver line, the first generation (G0) of injected adults were separated based on sex and crossed to 5-fold wild-type counterparts. Their offspring (F1) were manually screened for ECFP-derived cyan eye fluorescence. Single cyan-eyed F1 male was backcrossed for five generations to five wild-type females in order to establish a stable transgenic reporter line which later would be used for crossing with the driver line.

### Blood-feeding assay

Blood-feeding and host preference (below) assays were carried out as described in Wang et al^72^ with minor modifications. The experimental subjects were wildtype and *Orco^DsRed/DsRed^* homozygous mutant *Ae. albopictus*. Each trial used 30-35 non-blood-fed, 3-day-old, mated adult females. These mosquitoes were starved for 12 hours prior to the experiment. A 20-minute blood-feeding assay was conducted using mice immobilized with adhesive tape and a funnel-shaped wire mesh cage, during which mosquitoes primarily fed on the mouse’s tail. The experiments were performed in 18 cm × 18 cm × 18 cm nylon mesh cages, with 3 replicates conducted for each mosquito strain. The number of blood-fed mosquitoes was determined by visual inspection as fed or unfed. The ratio of blood-fed mosquitoes was calculated using the following formula: blood-fed (%) = Nb/Nt, where Nb was the number of blood-fed mosquitoes and Nt was the total number of mosquitoes.

### Host preference assay

The host preference assay was performed on wildtype and *Orco^DsRed/DsRed^* homozygous mutant *Ae. albopictus*. Each trial used 30-35 non-blood-fed, 3-day-old, mated adult females, which were fasted for 24 h prior to assay. Before testing, the mosquitoes were transferred to a two-choice olfactometer (20 cm x 20 cm x 32 cm) and allowed to acclimate for 30 min. The two ports of the olfactometer were respectively equipped with a human hand and a mouse. The participating volunteer (a 27-year-old female) had refrained from using any scented products for 48 hours prior to the experiment and had not washed her hands for 4 hours. The test mouse was immobilized adhesive tape and a funnel-shaped wire mesh cage. Fans were used to direct odors from the human hand and mouse, respectively, into the olfactometer toward female mosquitoes at an airflow speed of 0.5 m/s. The experiment lasted for 10 minutes, after which the number of female mosquitoes captured in the trap was counted and recorded. Host preference was expressed as the preference index = (Nh – Nm)/(Nh + Nm), where Nh was the number of mosquitoes probing human hand and Nm the number of mosquitoes probing mice. Six replications for each assay were performed.

### Data analysis and statistics

All statistical analysis was done using Prism 5 (GraphPad Software). Data are presented as mean ± s.e.m. Unpaired Student’s *t*-tests was used to compare two sets of data. The significance for all the tests was set to a P-value <0.05.

## Acknowledgement

This work was supported by the National Natural Science Foundation of China under Grants 82372289 and 82350410493 awarded to F.L. and S.T.F., respectively, and by Guizhou Education Department (Grant No. Qian Jiao Ji [2024] 54) and Guizhou Normal University (Grant No. QSXM [2022] B09) awarded to Z.Y., and the China Postdoctoral Science Foundation (2024M752152) to Q.Q..

## Authors’ contributions

Conceived and designed the study: FL, ZY, HY, QQ. Performed the experiments: HY, QQ, DG, HZ, FL. Prepared the materials: FL, ZY, SL. Wrote the paper: HY, QQ, ZY, STF, FL.

---

**Supplementary Table 1.**
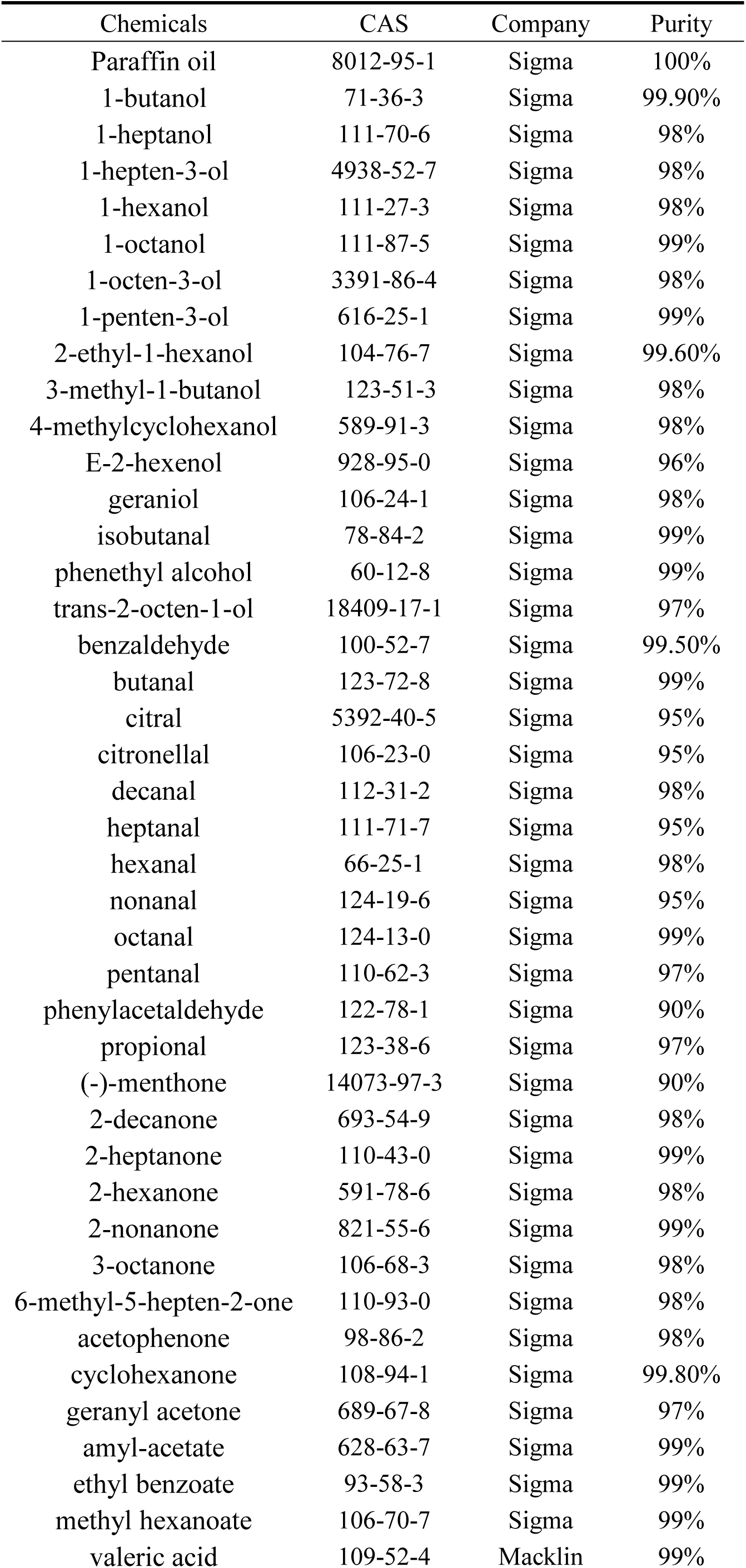
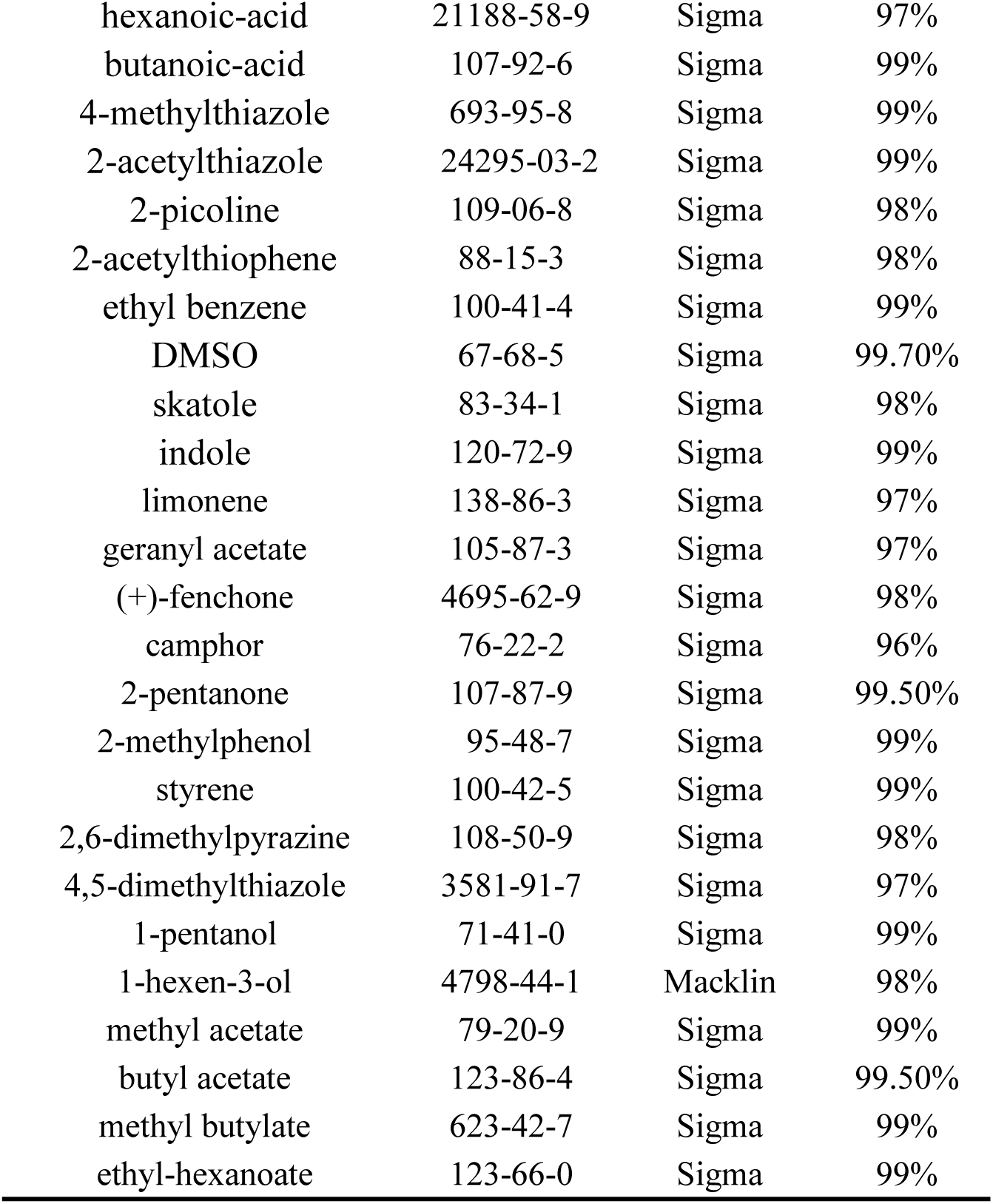
Odorants lists used in electrophysiological recordings.

