## Supplemental Table 1 for "Unveiling the Developmental Dynamics and Functional Role of Odorant Receptor Co-receptor (*Orco*) in *Aedes albopictus*: A Novel Mechanism for Regulating Tuning Odorant Receptor Expression"

Supplementary Table 1. Odorants lists used in electrophysiological recordings

| Chemicals | CAS | Company | Purity |
|---|---|---|---|
| Paraffin oil | 8012-95-1 | Sigma | 100% |
| 1-butanol | 71-36-3 | Sigma | 99.90% |
| 1-heptanol | 111-70-6 | Sigma | 98% |
| 1-hepten-3-ol | 4938-52-7 | Sigma | 98% |
| 1-hexanol | 111-27-3 | Sigma | 98% |
| 1-octanol | 111-87-5 | Sigma | 99% |
| 1-octen-3-ol | 3391-86-4 | Sigma | 98% |
| 1-penten-3-ol | 616-25-1 | Sigma | 99% |
| 2-ethyl-1-hexanol | 104-76-7 | Sigma | 99.60% |
| 3-methyl-1-butanol | 123-51-3 | Sigma | 98% |
| 4-methylcyclohexanol | 589-91-3 | Sigma | 98% |
| E-2-hexenol | 928-95-0 | Sigma | 96% |
| geraniol | 106-24-1 | Sigma | 98% |
| isobutanal | 78-84-2 | Sigma | 99% |
| phenethyl alcohol | 60-12-8 | Sigma | 99% |
| trans-2-octen-1-ol | 18409-17-1 | Sigma | 97% |
| benzaldehyde | 100-52-7 | Sigma | 99.50% |
| butanal | 123-72-8 | Sigma | 99% |
| citral | 5392-40-5 | Sigma | 95% |
| citronellal | 106-23-0 | Sigma | 95% |
| decanal | 112-31-2 | Sigma | 98% |
| heptanal | 111-71-7 | Sigma | 95% |
| hexanal | 66-25-1 | Sigma | 98% |
| nonanal | 124-19-6 | Sigma | 95% |
| octanal | 124-13-0 | Sigma | 99% |
| pentanal | 110-62-3 | Sigma | 97% |
| phenylacetaldehyde | 122-78-1 | Sigma | 90% |
| propional | 123-38-6 | Sigma | 97% |
| (-)-menthone | 14073-97-3 | Sigma | 90% |
| 2-decanone | 693-54-9 | Sigma | 98% |
| 2-heptanone | 110-43-0 | Sigma | 99% |
| 2-hexanone | 591-78-6 | Sigma | 98% |
| 2-nonanone | 821-55-6 | Sigma | 99% |
| 3-octanone | 106-68-3 | Sigma | 98% |
| 6-methyl-5-hepten-2-one | 110-93-0 | Sigma | 98% |
| acetophenone | 98-86-2 | Sigma | 98% |
| cyclohexanone | 108-94-1 | Sigma | 99.80% |
| geranyl acetone | 689-67-8 | Sigma | 97% |
| amyl-acetate | 628-63-7 | Sigma | 99% |
| ethyl benzoate | 93-58-3 | Sigma | 99% |
| methyl hexanoate | 106-70-7 | Sigma | 99% |
| valeric acid | 109-52-4 | Macklin | 99% |

| | | | |
|---|---|---|---|
| hexanoic-acid | 21188-58-9 | Sigma | 97% |
| butanoic-acid | 107-92-6 | Sigma | 99% |
| 4-methylthiazole | 693-95-8 | Sigma | 99% |
| 2-acetylthiazole | 24295-03-2 | Sigma | 99% |
| 2-picoline | 109-06-8 | Sigma | 98% |
| 2-acetylthiophene | 88-15-3 | Sigma | 98% |
| ethyl benzene | 100-41-4 | Sigma | 99% |
| DMSO | 67-68-5 | Sigma | 99.70% |
| skatole | 83-34-1 | Sigma | 98% |
| indole | 120-72-9 | Sigma | 99% |
| limonene | 138-86-3 | Sigma | 97% |
| geranyl acetate | 105-87-3 | Sigma | 97% |
| (+)-fenchone | 4695-62-9 | Sigma | 98% |
| camphor | 76-22-2 | Sigma | 96% |
| 2-pentanone | 107-87-9 | Sigma | 99.50% |
| 2-methylphenol | 95-48-7 | Sigma | 99% |
| styrene | 100-42-5 | Sigma | 99% |
| 2,6-dimethylpyrazine | 108-50-9 | Sigma | 98% |
| 4,5-dimethylthiazole | 3581-91-7 | Sigma | 97% |
| 1-pentanol | 71-41-0 | Sigma | 99% |
| 1-hexen-3-ol | 4798-44-1 | Macklin | 98% |
| methyl acetate | 79-20-9 | Sigma | 99% |
| butyl acetate | 123-86-4 | Sigma | 99.50% |
| methyl butylate | 623-42-7 | Sigma | 99% |
| ethyl-hexanoate | 123-66-0 | Sigma | 99% |
